# Molecular identification of ALDH1A1 and SIRT2 in the astrocytic putrescine-to-GABA metabolic pathway

**DOI:** 10.1101/2023.01.11.523573

**Authors:** Mridula Bhalla, Jeong Im Shin, Yeon Ha Ju, Yongmin Mason Park, Seonguk Yoo, Hyeon Beom Lee, C Justin Lee

## Abstract

GABA (γ-aminobutyric acid) is the primary inhibitory neurotransmitter in the CNS. In astrocytes, GABA is synthesized by degradation of putrescine by monoamine oxidase B (MAO-B), a process which is known to mediate tonic inhibition of neuronal excitability. This astrocytic tonic GABA and related enzymes are also reported to be involved in memory impairment in Alzheimer’s Disease, and therefore are potential therapeutic targets to rescue memory in AD patients. However, the enzymes downstream of MAO-B in this pathway have not been elucidated yet. To fill this gap in knowledge, we performed transcriptomic and literature database analysis and identified Aldehyde dehydrogenase 1 family, member A1 (ALDH1A1) and a histone deacetylase enzyme Sirtuin2 (SIRT2) as plausible candidate enzymes in primary cultured astrocytes. Immunostaining, metabolite analyses, and sniffer patch clamp performed in the presence or absence of suitable inhibitors, or with genetic ablation of the candidate enzymes recapitulated their participation in GABA production. We propose ALDH1A1 and SIRT2 as potential therapeutic targets against Alzheimer’s Disease.

## INTRODUCTION

Astrocytes in the brain have been acknowledged to play a role in maintaining homeostasis at the synapse, regulating neuronal signaling and protecting neurons from oxidative damage (Chen et al., 2020). Their role in the production and tonic release of the inhibitory neurotransmitter γ-aminobutyric acid (GABA) in neurodegenerative diseases such as Alzheimer’s Disease (AD) and Parkinson’s Disease (PD) has been highlighted recently (Heo et al., 2020; Jo et al., 2014; Nam et al., 2020; Woo et al., 2018). However, previous studies and our current knowledge of GABA production in astrocytes only provide a partial picture about the molecular players involved in the process, particularly the participating enzymes and their regulatory roles. A comprehensive understanding of this process can aid us to better realize therapeutic strategies against neurodegenerative disorders and target GABA production in a more disease-specific manner.

GABA in astrocytes can have two major sources: the GABA that is taken up from extracellular spaces by the GABA transporter, and the endogenous synthesis of GABA from precursors glutamate or putrescine (Ishibashi et al., 2019). Glutamic acid decarboxylase (GAD65 or GAD67)-mediated conversion of glutamate to GABA has been hypothesized but the causal relationship between the expression of GAD65/67 in astrocytes and GABA synthesis has not been clearly determined in previous studies (Lee et al., 2011). Putrescine is known to be converted to GABA via monoamine oxidase B (MAOB)-dependent degradation (Yoon et al., 2014). In addition to the MAOB-dependent pathway, the existence of an alternate diamine oxidase (DAO)-mediated conversion of putrescine to GABA has also been elucidated in thalamic and hippocampal astrocytes (Kwak et al., 2020; Park et al., 2019).

MAOB-mediated conversion of putrescine to GABA is a 4-step pathway (Park et al., 2019; Yoon et al., 2014), as opposed to the 2-step conversion mediated by DAO and ALDH1A1 (Kwak et al., 2020). Putrescine is converted to N-acetyl-putrescine in the presence of Coenzyme A by enzyme putrescine acetyl transferase (PAT; also known as spermine/spermidine acetyl transferase SSAT1/SAT1) (Heo et al., 2020), which is further oxidized to N-acetyl-γ-aminobutyraldehyde by MAOB (Yoon et al., 2014). The enzymes downstream to MAOB in this pathway have not been well studied, although their enzymatic functions can be easily predicted based on the intermediate metabolites in the pathway (Fig. 1A). Following oxidation by MAOB, the intermediate aldehyde is further oxidized to N-acetyl-GABA by an aldehyde dehydrogenase (ALDH) family member, currently speculated to be ALDH2 (Yoon and Lee, 2014) and widely known for its role in alcohol metabolism in the liver (Edenberg, 2007). Researchers have shown the involvement of ALDH2 in monoamine metabolism in mitochondrial extracts from rat livers (Keung and Vallee, 1998) and in the metabolism of ethanol to produce GABA in cerebellar astrocytes (Jin et al., 2021), but there is no report of its involvement in the production of GABA from putrescine. However, among the 19 known ALDH family members, it has been reported that ALDH 1 family member A1 (ALDH1A1) mediates the synthesis of GABA in midbrain dopaminergic neurons (Kim et al., 2015) and thalamic astrocytes (Kwak et al., 2020), and is highly expressed in adult mouse hippocampal astrocytes (Chai et al., 2017), raising the possibility of the involvement of ALDH1A1 in astrocytic MAOB-mediated conversion of putrescine to GABA as well.

**Figure 1.**
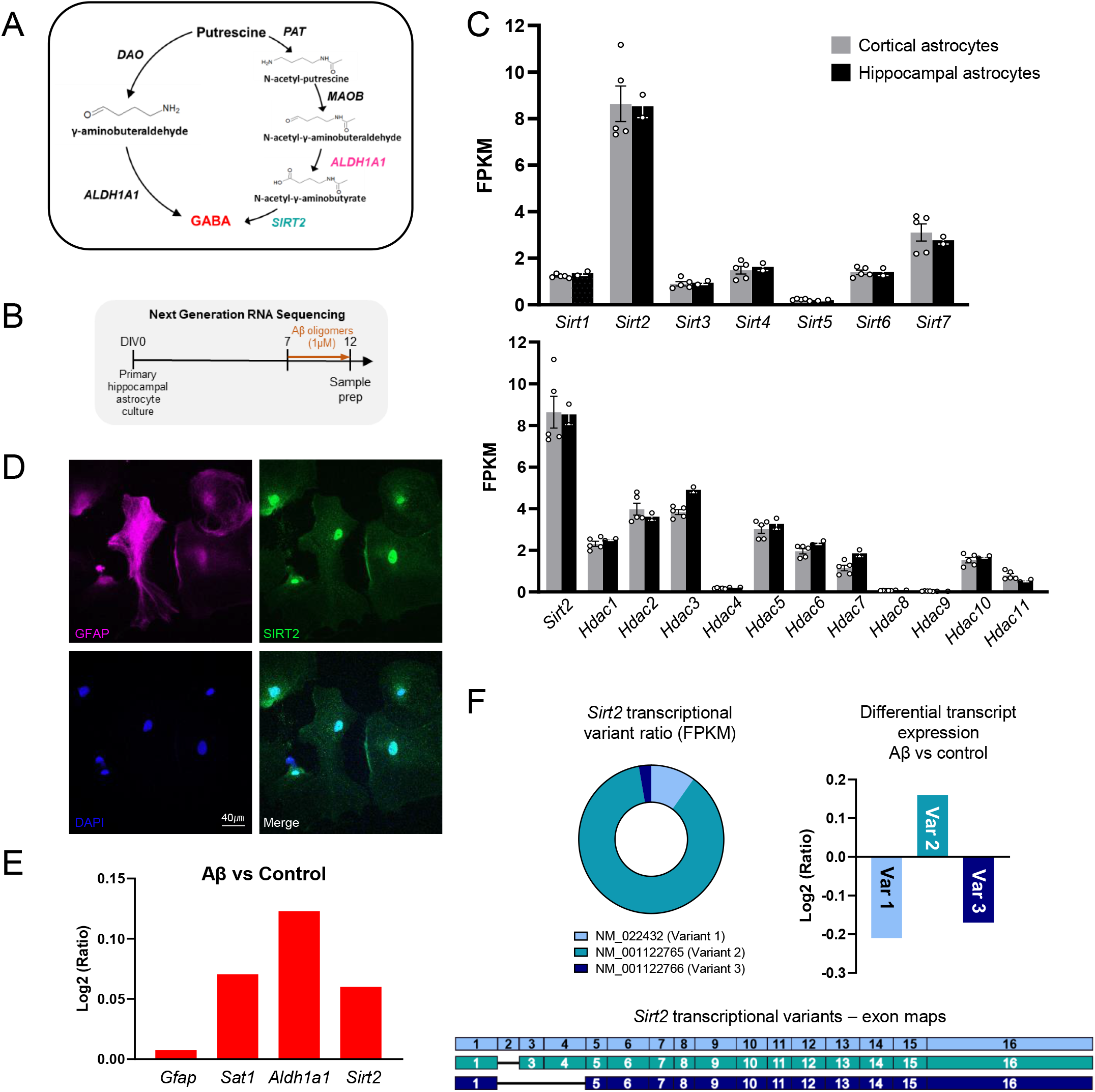
*Sirt2* is highly expressed in cortical and hippocampal astrocytes. (A) Schematic diagram of the putrescine-to-GABA conversion pathways with predicted candidate enzymes. (B) Experimental timeline for Next Generation RNASeq. (C) Bar graph of FPKM values of sirtuin family members (top) and histone deacetylases (bottom) in primary cortical and hippocampal astrocyte cultures. (D) Immunostaining for SIRT2 and GFAP in primary astrocyte cultures. (E) Differential expression analysis of genes using RNASeq in astrocytes upon Aβ treatment. (F) Top Left, pie chart representation of ratio of *Sirt2* transcriptional variants in astrocytes; Top Right, bar graph of differential expression analysis of *Sirt2* transcriptional variants; Bottom, representation of exon map of *Sirt2* transcriptional variants. Data represents Mean ± SEM.

Further, the N-acetyl-GABA is deacetylated by an unknown deacetylase to finally synthesize GABA (Le-Corronc et al., 2011). Major protein deacetylases in the cell can be classified as histone deacetylases (HDACs) or sirtuins (SIRTs). There is mounting evidence highlighting the role of sirtuins in several models of neurodegeneration. There are 7 proteins belonging to the SIRT family in humans, all of which have strikingly been implicated in neurodegenerative disorders (Yeong et al., 2020). These sirtuins, although known to primarily be histone deacetylases, have also been found to be localized in non-neuronal subcellular regions, thereby participating in other cellular pathways and processes. Among the 7 sirtuin proteins, the beneficial effect of SIRT2 inhibition in Alzheimer’s Disease, Parkinson’s Disease and Huntington’s Disease (Yeong et al., 2020) indicates its potential role in neurodegeneration and disease pathogenesis. We therefore hypothesize that SIRT2 participates in the deacetylation of N-acetyl-GABA to GABA in astrocytes.

In this study, we sought to determine the enzymes downstream to MAOB involved in the astrocytic putrescine-to-GABA conversion pathway. We hypothesized ALDH1A1 and sirtuins to be potential candidates for this, due to their notable expression levels and participation in astrocytic GABA-associated disease pathology. To investigate our hypothesis, we performed Next Generation RNA-sequencing (NGS) to detect the expression of different deacetylases in astrocytes and changes in their levels in AD-like conditions. In cultured astrocytes, we measured the changes in putrescine-induced GABA and N-acetyl-GABA levels on pharmacological and genetic manipulation of SIRT2 activity. We also measured the production and release of GABA from cultured astrocytes on pharmacological and genetic manipulation of ALDH1A1 as well as SIRT2 to confirm our findings. Indeed, we were able to demonstrate that SIRT2 and ALDH1A1 were majorly involved in the putrescine-induced production of GABA in astrocytes.

## METHODS

### Primary astrocyte culture

Primary astrocytes were cultured from P1 pups of C57BL/6J mice as previously described (Woo et al., 2012). Briefly, the cerebral cortex and hippocampus were dissected and cleaned of meninges and midbrain before dissociation into a single cell suspension by trituration in astrocyte culture medium. The medium was prepared by using Dulbeccos’ modified Eagle’s Medium (DMEM, Corning) supplemented with 4.5 g/L glucose, L-glutamine, sodium pyruvate, 10% heat-inactivated horse serum, 1% heat-inactivated fetal bovine serum and 1000 units/mL of penicillin-streptomycin. Cells were plated onto culture dishes coated with 0.1mg/mL poly-D-lysine (Sigma) and maintained in astrocyte culture medium at 37°C in a humidified atmosphere containing 5% CO_2_. Three days later (at DIV4), cells were vigorously washed with Dulbecco’s phosphate buffered saline by repeated pipetting and the media was replaced.

### Illumina Hiseq library preparation and RNA sequencing

RNA was isolated from cultures primary astrocytes using Qiagen RNEasy Kit (Qiagen, #74104). Sample libraries were prepared using the Ultra RNA Library Prepkit (NEBNext, #E7530), Multiplex Oligos for Illumina (NEBNext, #E7335) and polyA mRNA magnetic isolation module (Invitrogen, #61011) following manufacturers’ instructions. Full details of the library preparation and sequencing protocol are provided on the website and previously described (Ju et al., 2022). The Agilent Bioanalyser and associated High Sensitivity DNA Kit (Agilent Technologies) were used to determine the quality, concentration, and average fragment length of the libraries. The sample libraries were prepared for sequencing according to the HiSeq Reagent Kit Preparation Guide (Illumina, San Diego, CA, USA). Briefly, the libraries were combined and diluted to 2nM, denatured using 0.1N NaOH, diluted to 20pM by addition of Illumina HT1 buffer and loaded into the machine along with read 1, read 2 and index sequencing primers. After the 2×100 bp (225 cycles) Illumina HiSeq paired-end sequencing run was complete, the data were base called and reads with the same index barcode were collected and assigned to the corresponding sample on the instrument, which generated FASTQ files for analysis.

### NGS Data Analysis

BCL files obtained from Illumina HiSeq2500 were converted to fastq and demultiplexed based on the index primer sequences. The data was imported to Partek Genomics Suite (Flow ver 10.0.21.0328; copyright 2009, Partek, St Louis, MO, USA), where the reads were further processed. Read quality was checked for the samples using FastQC. High quality reads were aligned to the *Mus musculus* (mouse) genome assembly GRCm38 (mm10, NCBI) using STAR (2.7.3a). Aligned reads were quantified to the mouse genome assembly (mm10, RefSeq transcripts 93) and normalized to obtain fragments per kilobase million (or FPKM) values of positively detected and quantified genes. Gene read counts were also normalized to Transcripts per million (TPM), which was used to identify alternate splice variants of the positively detected genes. Differential gene analysis was carried out by normalizing the quantified and annotated gene reads to the Median Ratio and performing DeSeq2 (available on Partek Genomics Suite).

### Immunocytochemistry

For pharmacological study, astrocytes (DIV 7-10) were seeded on coverslips and incubated with 180mM Putrescine in the presence or absence of 10uM DEAB, 200nM EX527 or 3uM AGK2 overnight. For genetic ablation study, DIV 7 astrocytes were detached from culture dish surface, electroporated with mCherry-tagged pSicoR vector carrying Scr sequence or shRNA sequences against SIRT2 or Aldh1a1 (shSIRT2 targeting 5’-GGAGCATGCCAACATAGATGC-3’ or shAldh1a1 targeting 5’-TTTCCCACCATTGAGTGCC-3’ respectively) and seeded onto coverslips. Two days later, they were treated with 180uM putrescine for 24 hours. Cells on the coverslips were fixed with 4% paraformaldehyde (Sigma-Aldrich) in 0.1M PBS at room temperature for 15 minutes. After fixation, the coverslips were washed 3 times with 0.1M PBS for 10 minutes each, then blocked with 0.1M PBS containing 0.3% Triton X-100 (Sigma, USA) and 10% Donkey Serum (Genetex) for 1.5 hrs at room temperature. The cells were then incubated with primary antibodies in blocking solution in the following composition: guinea-pig anti-GABA antibody (1:1000, AB175, Millipore, USA), chicken anti-GFAP antibody (1:1000, AB5541, Millipore, USA) for overnight (atleast 16 hours) at 4°C with gentle rocking. After washing 3 times with 0.1M PBS, 10 minutes each, the cells were incubated with corresponding secondary antibodies in blocking solution in the following composition: conjugated Alexa 594 chicken anti IgG (1:500, 703-585-155, Jackson, USA) or Alexa 647 donkey anti-guinea-pig IgG (1:500, 706-605-148, Jackson, USA) for 2 hours at room temperature with gentle rocking. The cells were then incubated with 1:2000 DAPI solution (Pierce) in 0.1M PBS for 10 minutes followed by 3 rinses with 0.1M PBS. Cover slips were finally mounted onto slide glass with fluorescence mounting solution (S3023, DAKO, USA). Images were acquired using a Nikon A1R confocal microscope (pharmacological study) or Zeiss LSM900 microscope (genetic ablation study) and analysed using the ImageJ program (NIH).

### Metabolite analysis

For metabolite analysis, electrospray ionization LC-MS/MS was used. Exion LC™ AD UPLC which was coupled with an MS/MS (Triple Quad 4500 System, AB Sciex LLC, Framingham, USA) using an Acquity^®^ UPLC BEH HILIC column (1.7 etyl-GABA, and GA 2.1 mm × 100 mm, Waters, USA) at 30°C, has been used and the system was controlled by Analyst 1.6.2 software (AB Sciex LP, Ontario, Canada). 70% methanol was added to the astrocyte sample pellets and the mixture was vortexed for 30s. The lysate from the cells, which was produced by three consecutive freeze-thaw cycles using liquid nitrogen, was centrifuged for 10 minutes at ~21000g (14,000rpm). 5μL of supernatant from each sample was used for DNA normalization (Nano-MD UV-Vis spectrophotometer; Scinco, Seoul). 40μL of the supernatant from each sample was evaporated to dryness at 37°C under a gentle stream of nitrogen. Phenylisothiocyanate (PITC) derivatization was performed by adding 50μL of mixture of ethanol, water, pyridine and PITC (19:19:19:3 v/v), vortexing for 30s and shaking for 20 min, followed by evaporating to dryness at 37°C under a gentle stream of nitrogen. The residue was reconstituted by adding 50μL of the mobile phase A (0.2% formic acid in deionized water): B (0.2% formic acid in acetonitrile) = 5:5 solvent and vortexing for 30s. The initial chromatographic conditions were 100% solvent A at a flow rate of 0.4 mL·min-1. After 0.9min at 15% B, solvent B was set to 15% over the next 4.1min, solvent B was set to 70% over the next 5min, solvent B was set to 100% over the next 0.5min, and these conditions were retained for an additional 2min. The system was then returned to the initial conditions over the next 0.5min. The system was re-equilibrated for the next 2.5min in the initial conditions. The total running time was 15min. All samples were maintained at 4°C during the analysis, and the injection volume was 5μL. The MS analysis was performed using ESI in positive mode. The ion spray voltage and vaporizer temperature were 5.5 kV and 500°C, respectively. The curtain gas was kept at 45 psi, and the collision gas was maintained at 9 psi. The nebulizer gas was 60 psi, while the turbo gas flow rate was 70 psi. The metabolites were detected selectively using their unique multiple reaction monitoring (MRM) pairs. The following MRM mode (Q1 / Q3) was selected: putrescine (m/z 359.200 / 266.100), GABA (m/z 238.875/ 87.103). As to monitor specific parent-to-product transitions, the standard calibration curve for each metabolite was used for absolute quantification.

### 2-cell sniffer patch clamp recording

Primary astrocyte cultures were prepared from P1 C57BL/6 mouse pups as described above. As required, the cells were seeded onto poly-D-lysine-coated cover glass and either electroporated with respective shRNA constructs (genetic ablation experiments) or treated with inhibitors in the presence of putrescine (pharmacological inhibition) on DIV7. On the day of sniffer patch, HEK 293-T cells expressing GFP-tagged GABA_C_ receptors were seeded onto the astrocytes and allowed to settle for atleast 1 hour before patching. The cover glasses were then immersed in 5μM Fura-2-AM (in 1mL external HEPES solution containing 5μL of 20% pluronic acid) for 40 minutes to allow Fura incorporation into the cell, washed at room temperature (with external solution, described later) and subsequently transferred to the microscope stage. The external solution of following composition (in mM): 150 NaCl, 10 HEPES, 3 KCl, 2 CaCl_2_, 2 MgCl_2_, 5.5 glucose (pH adjusted to 7.3, osmolality to 320 mOsmol kg^-1^) was allowed to continuously flow over the cells during the experiment, and during full activation recording, was replaced with one containing 100μM GABA. Images at 510nm wavelength were taken after excitation by 340nm and 380nm light using pE-340^fura^ (CoolLED) to record calcium transients within the cells. The two resulting images were used for ratio calculations in Axon Imaging Workbench (version 11.3, Axon Instruments). To perform sniffer patch, the astrocytic TRPA1 receptor was activated by pressure poking with a glass pipette and the resulting GABA release was recorded as inward current in the GABAC-expressing HEK 293T cells under voltage clamp (V_h_ = −60mV) using Axopatch 200A amplifier (Axon Instruments), acquired with pClamp 11.3. Recording electrodes (4-10 MΩ) were filled with the following internal solution (in mM): 110 Cs-gluconate, 30 CsCl, 0.5 CaCl_2_, 10 HEPES, 4 Mg-ATP, 0.3 Na3-GTP and 10 BAPTA (pH adjusted to 7.3 with CsOH, osmolality adjusted to 300mOsm kg^-1^ with sucrose). For simultaneous recording of calcium response with the patch and poking pipettes, Imaging Workbench was synchronized with pClamp 11.3. To account for differences in GABA_C_ receptor expression on the HEK cells, saturating concentration of 100μM GABA (in HEPES solution) was applied to record maximal GABA current from the cell, and the poking-induced current was normalized as percentage of full activation current on application of GABA from the HEK cell.

## RESULTS

### Next Generation Sequencing reveals high expression of SIRT2 in primary cultured astrocytes

To begin, we examined the putrescine-to-GABA conversion pathway in the brain and dissected the molecular processes involved in each step (Fig. 1A). To investigate the presence of deacetylases in primary cultured astrocytes, we performed Next Generation RNA-Sequencing (NGS) to objectively compare the expression levels of our candidate deacetylases (Fig. 1B) in primary astrocyte cultures derived from cortex as well as hippocampus of P1 mice. We screened the FPKM levels of different Sirtuin genes and histone deacetylases and found that Sirtuin 2 (*Sirt2*) was expressed at the highest level in both cortical as well as hippocampal astrocytes (Fig. 1C). Although SIRT2 is majorly known for its role in microtubule deacetylation, the protein can also translocate to the nucleus to modulate the cell cycle (Grabowska et al., 2017). Using immunocytochemistry, we determined the localization of the SIRT2 protein in the cytoplasm as well as the nucleus of astrocytes (Fig. 1D), indicating that it actively participates in cellular metabolic processes in astrocytes. We found that 5-day treatment of Aβ oligomers (1μM), which has been known to induce Alzheimer’s Disease-like astrocyte reactivity *in vitro* (Ju et al., 2022) upregulated the expression of candidate enzymes involved in the putrescine-to-GABA degradation pathway (Fig. 1E). Furthermore, transcriptional variant analysis shows that transcriptional variant 2 of mouse *Sirt2* (NM_001122765.2), which is the majorly expressed variant in the central nervous system that translates to a functional protein (Maxwell et al., 2011), was specifically upregulated (Fig. 1F), indicating that SIRT2 was a reasonable candidate for our proposed deacetylase enzyme.

### SIRT2, not SIRT1, and ALDH1A1 are involved in the conversion of putrescine to GABA in primary cultured astrocytes

To determine the extent of GABA production by putrescine treatment in cultured astrocytes, we treated primary hippocampal astrocyte culture with 180μM putrescine for 1 day and performed immunocytochemistry (ICC) for GFAP and GABA to compare directly against 100μM GABA treatment (Fig. 2A-C). Upon analysis, we found that 1-day treatment of putrescine was sufficient to increase GABA production in the astrocytes up to half that of GABA-treated cultures (734.7 ± 44.45 a.u. vs. 1386 ± 198.6 a.u. respectively; Fig. 2C). To confirm the role of SIRT2 in GABA synthesis, we co-treated the cultures with putrescine and inhibitors for SIRT1 and SIRT2, EX527 (200nM) and AGK2 (3μM) respectively (Fig. 2D). While EX527 treatment had no effect on putrescine-induced GABA levels in the cells, we saw a significant reduction in GABA staining in the cells treated with SIRT2 inhibitor AGK2 (Fig. 2E), confirming that SIRT2 was involved in the GABA-production pathway. To further rule out the inhibition of any non-specific deacetylase by AGK2, we synthesized silencing hairloop RNA (shRNA) sequence specific to *Sirt2* in electroporated the astrocytes to genetically knockdown the expression of SIRT2. Additionally, to check for a potential aldehyde dehydrogenase candidate enzyme involved in GABA production, we electroporated the cultured astrocytes with shRNA specific to *Aldh1a1* (Kwak et al., 2020) and checked GABA levels after 1 day of putrescine treatment (Fig. 2F). As a result, putrescine-induced GABA levels were reduced in primary cultured astrocytes upon genetic ablation of *Aldh1a1* as well as *Sirt2* (Fig. 2G), suggesting their role in putrescine-to-GABA conversion in astrocytes. Taken together, we suggest that ALDH1A1 and SIRT2 were the unknown enzymes downstream to MAOB in the GABA production pathway.

**Figure 2.**
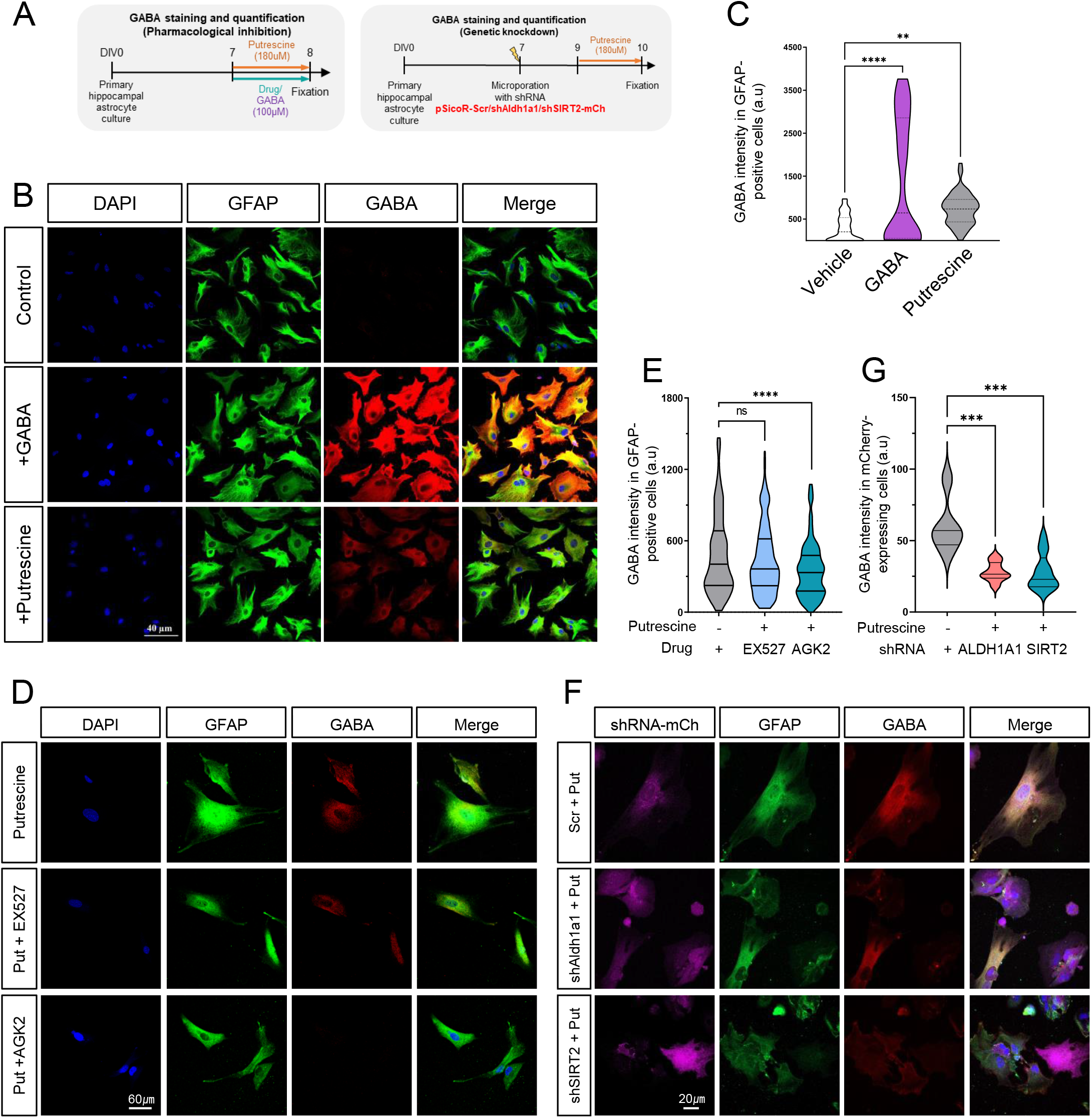
SIRT2, not SIRT1, and ALDH1A1 are involved in the conversion of putrescine to GABA in astrocytes. (A) Experimental timeline for GABA staining and quantification. (B, D, F) Immunostaining for GFAP and GABA in primary cultured astrocytes treated with putrescine or GABA (B), putrescine in the presence or absence of EX527 or AGK2 (D) or putrescine treated primary cultured astrocytes expressing Scr/shALDH1A1/shSIRT2-mCherry (F) (C, E, G) Truncated violin plot for GABA intensity in GFAP-positive cells from (B) and (D), and mCherry-positive (shRNA-expressing) cells in (F). Data represents Mean ± SEM. **, p<0.01; ***, p<0.001; ****, p<0.0001 (Ordinary one-way ANOVA)

### Intermediate metabolite in putrescine-to-GABA conversion accumulates on inhibition or genetic knockdown of SIRT2

To investigate the direct changes in the intermediates formed during the conversion to putrescine to GABA, we performed liquid chromatography-mass spectrometry (LC-MS) analysis in primary cultured astrocytes after 1 day treatment of putrescine in the presence and absence of SIRT2 inhibitor AGK2 (Fig. 3A and B). We found that intracellular levels of putrescine and SIRT2 substrate N-acetyl GABA were about 3-fold higher in putrescine treated cells and remained unchanged on SIRT2 inhibition (Fig. 3B). Consistent with our previous findings (Fig. 2), GABA levels increased on putrescine treatment, which were interestingly brought down to control levels upon inhibition of SIRT2. We further corroborated our results by genetically ablating SIRT2 via shRNA (Fig. 3A and C). While the data for putrescine levels was consistent with our pharmacological inhibition experiments, it was noteworthy that we observed over 1.5-fold accumulation of N-Acetyl-GABA upon knockdown of SIRT2 in putrescine-treated astrocytes, indicating its role in the deacetylation of the intermediate to form GABA. Intracellular GABA levels were similarly decreased by SIRT2 inhibition. These results indicate that SIRT2 is a key enzyme in the production of GABA from putrescine, via the deacetylation of the intermediate metabolite N-Acetyl-GABA.

**Figure 3.**
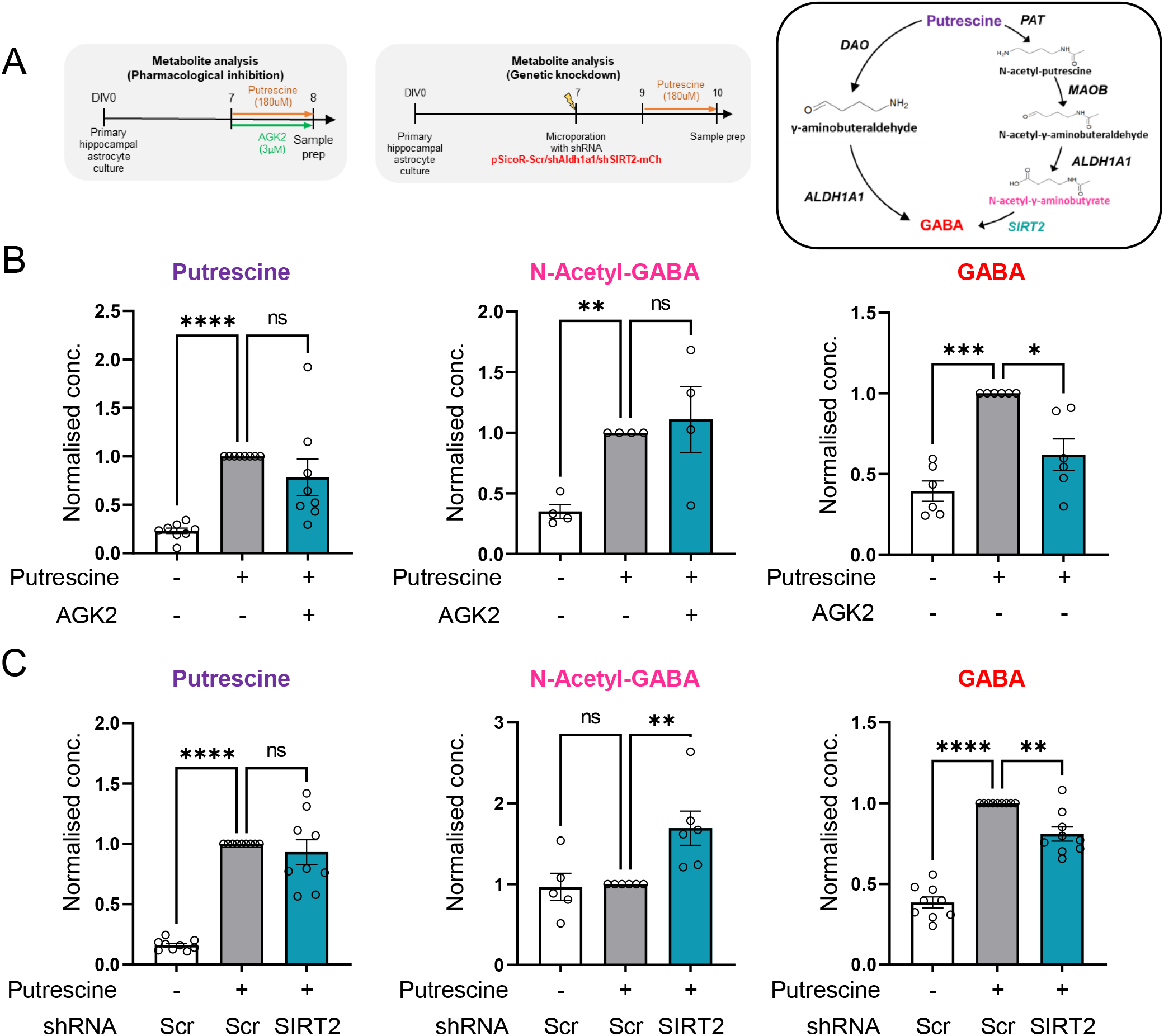
SIRT2 could be involved in the conversion of N-acetyl-GABA to GABA. (A) Experimental timeline for metabolite analysis experiments (Left, Middle) and schematic of putrescine-to-GABA conversion (Right) (B) Bar graphs for relative metabolite concentration in primary astrocyte cultures treated with or without putrescine, in the presence or absence of AGK2 (normalized to metabolite concentration in putrescine-treated cultures) (C) Bar graphs for relative metabolite concentration in primary cultures astrocytes expressing Scr/shSIRT2-mCherry treated with or without putrescine (normalized to metabolite concentration in putrescine-treated Scr-mCh-expressing culture) Data represents Mean ± SEM. *, p<0.05; **, p<0.01; ***, p<0.001; ****, p<0.0001 (RM one-way ANOVA)

### SIRT2 is essential, while ALDH1A1 is only partially responsible for GABA production from putrescine in astrocytes

To further examine the release of the GABA produced on accumulation of excess putrescine in astrocyte cultures, we performed 2-cell sniffer patch experiments on putrescine-treated astrocytes in the presence or absence of appropriate inhibitors (Fig. 4A and B). Briefly, on poking an astrocyte membrane, TRPA1 channels activate and cause an influx of calcium ions into the cell, which can be recorded as a calcium signal using fura-2-AM (Oh et al., 2020). This calcium influx causes GABA release from the cell via astrocytic BEST1 channel, which can be measured as current recorded on a nearby GABAC receptor-expressing HEK293T cell (Fig. 3A) and can be used as a measure of intracellular GABA level in the poked astrocyte. 1-day treatment of putrescine significantly increased the amount of GABA released from the astrocyte (Fig. 3C), which was substantially eliminated by the inhibition of ALDH1A1, using DEAB (N,N-diethylaminobenzaldehyde, 1μM), or SIRT2, using AGK2 (3μM) (Fig. 3D). As DEAB does not specifically inhibit ALDH1A1 (Morgan et al., 2015), we validated our results by using shRNA specific for *Aldh1a1* (Fig. 3E and F). Interestingly, we found that while DEAB was able to eliminate about 90% of GABA current when compared to vehicle treatment, shALDH1A1 was only able to eliminate 65% of the poking-induced GABA release, implying the role of other DEAB-sensitive aldehyde dehydrogenase enzymes in the conversion of putrescine to GABA.

**Figure 4.**
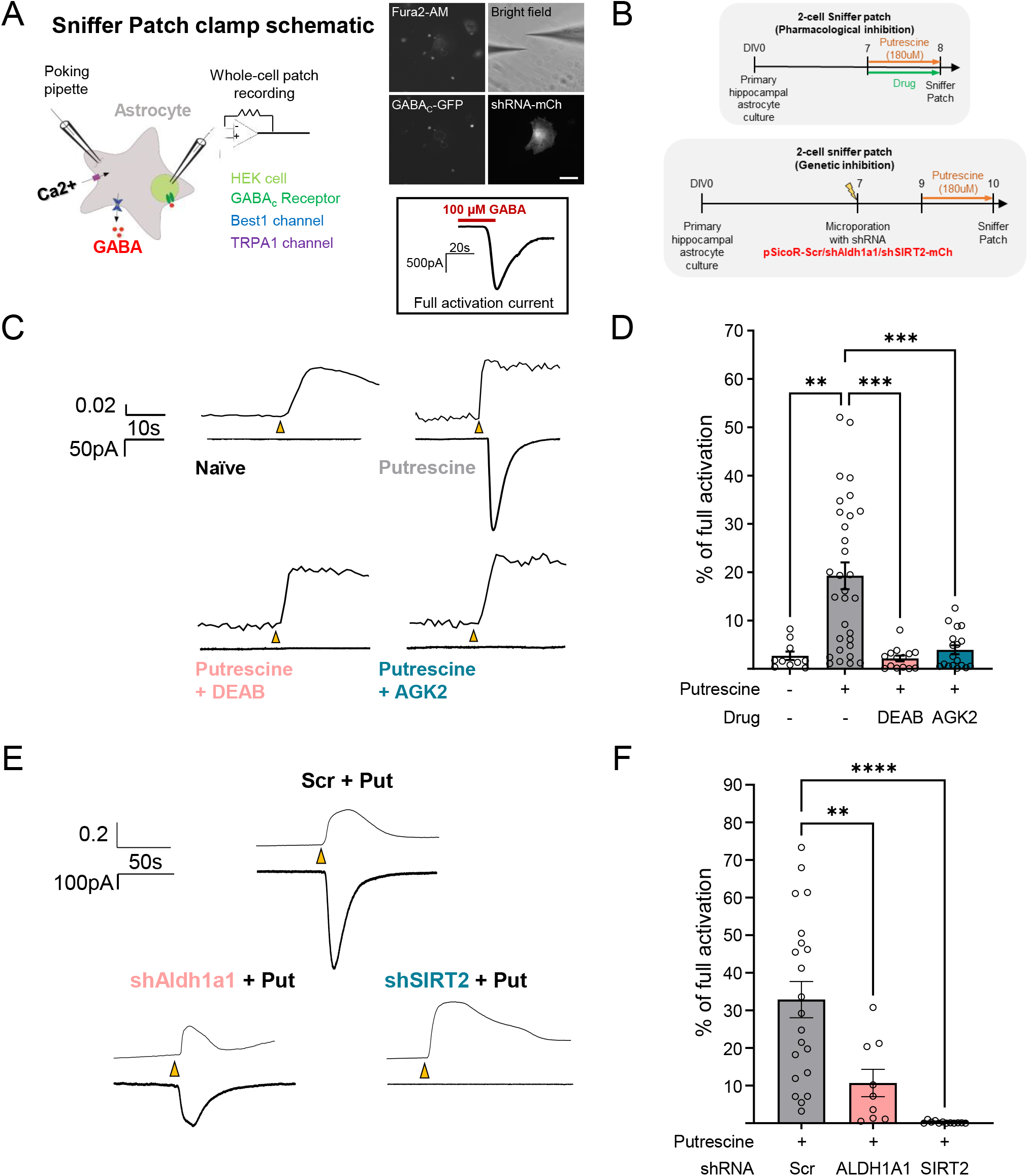
SIRT2 is essential, while ALDH1A1 is partially involved in GABA production. (A) Schematic for working of 2-cell sniffer patch experiments (Left) Representative fluorescence images of the sniffer patch experiment (Right top, scale bar 40μm) and representative trace of full activation of GABAC-expressing HEK cells by GABA treatment (Right bottom). (B) Experimental timeline for 2-cell sniffer patch experiments. (C, D) Representative traces (C) and bar graph (D) of GABA release-mediated sensor current from astrocytes treated with or without putrescine in the presence or absence of DEAB or AGK2 (E, F) Representative traces € and bar graph (F) of GABA release-mediated sensor current from putrescine-treated cultured astrocytes expressing Scr/shALDH1A1/shSIRT2-mCherry Data represents Mean ±SEM. *, p<0.05; **, p<0.01; ***, p<0.001; ****, p<0.0001 (Ordinary One-way ANOVA)

Genetic ablation of SIRT2 using shRNA was able to eliminate poking-induced GABA release from astrocytes (Fig. 3F). Taken together, our results indicate a partial role of ALDH1A1 and key role of SIRT2 in the production of GABA from putrescine in astrocytes.

## DISCUSSION

In this study, we have attempted to delineate the enzymes involved in the metabolism of putrescine to GABA mediated by MAOB. Based on RNASeq data analysis, we have identified SIRT2 as the best candidate for the final deacetylation step and demonstrated its role by pharmacological inhibition (AGK2) or gene silencing (shSIRT2) *in vitro* primary-cultured mouse astrocytes. Inhibition or genetic ablation of SIRT2 leads to reduced GABA production in primary cultured astrocytes on immunostaining (Fig. 2), metabolite analysis (Fig. 3) and 2-cell sniffer patch (Fig. 4). Furthermore, we also see accumulation of predicted SIRT2 substrate N-acetyl-GABA (Fig. 3), supporting our hypothesis. We reveal that ALDH1A1 also participates in putrescine-to-GABA conversion, as can be observed in Figures 2 and 4. Based on our findings, we propose that ALDH1A1 and SIRT2 are the enzymes downstream to MAOB in astrocytic GABA production pathway.

Our study provides the first line of evidence that SIRT2 is involved in GABA production in astrocytes. SIRT2 is majorly known for its role in the cell cycle via α-tubulin deacetylation (Li et al., 2007) and H4K16 deacetylation (Vaquero et al., 2006), and it has been reported that the levels of the protein increase as cells senesce (Grabowska et al., 2017). While it has previously been demonstrated that the inhibition of SIRT2 reduces astrocyte reactivity markers (Scuderi et al., 2014) and rescues α-synuclein-mediated toxicity in Parkinson’s Disease models (Outeiro et al., 2007), our study is the first to not only reveal the role of SIRT2 in astrocytic GABA production, but also isolate the metabolic step catalyzed by SIRT2. By analyzing the levels of intermediate metabolite N-acetyl-GABA in primary cultured astrocytes on inhibition or genetic ablation of the enzyme (Fig. 3), we suggest that SIRT2 is the deacetylase enzyme participating in the final step of the putrescine-to-GABA conversion. This must, however, be validated by testing the NAD+-dependency of this process, as SIRT2 is a NAD+-dependent deacetylase. Additionally, due to the role of SIRT2 in oligodendrocyte precursor cell proliferation and differentiation (Li et al., 2007), in-depth animal model studies are required to test the overall effect of SIRT2 inhibition in the brain on cognition and locomotion (Wang et al., 2019). While a potential therapeutic effect of SIRT2 inhibition against cervical cancer by cell cycle arrest has been reported (Singh et al., 2015), SIRT2 KO mice also show elevated rates of tumourigenesis (Wang et al., 2019), implying the crucial role of this protein in cell cycle regulation and thereby reducing the viability of SIRT2 inhibition as a therapeutic target against neurodegeneration. However, the discovery of the involvement of previously unexplored enzyme SIRT2 in GABA production leads to fascinating and exciting new prospects regarding its manipulation to specifically target astrocytic GABA production.

While ALDH2 has been reported to be involved in monoamine metabolism in the liver (Keung and Vallee, 1998) and alcohol metabolism in the brain (Jin et al., 2021), our NGS data reported upregulation of *Aldh1a1* in Aβ-treated AD-like astrocytes (Fig. 1), suggesting the involvement of ALDH-family members other than ALDH2 in this process. Putrescine-induced GABA production was inhibited by DEAB (Fig. 4), which is reported to have little, if any, turnover when incubated with ALDH2 (Morgan et al., 2015), further supporting that ALDH2 is not participating in this process. While the pharmacological inhibition of ALDH1A1 with DEAB was able to largely eliminate the putrescine-induced GABA release from cultured astrocytes (Fig. 4D), genetic knockdown only showed a partial elimination (Fig. 4F). As DEAB is well-known to be a non-specific inhibitor (Morgan et al., 2015), we predict that ALDH3A1, an ALDH-family member which uses DEAB as a substrate, could be involved in this oxidation step. There also exists the possibility that another ALDH-enzyme is switched-on in compensation of the genetic knockdown in Fig. 4E, F, and this possibility awaits further study.

Armed with knowledge of the molecular players involved in the production of inhibitory neurotransmitter GABA from astrocytes in neurodegenerative conditions, we can design better therapeutic strategies against these diseases. Identifying the partial role of ALDH1A1 is the first step towards this holistic approach and must be supplemented with more experiments towards demarcating the ALDH family members contributing to or participating in compensatory mechanisms involved in the oxidation step following MAOB. The identification of SIRT2 as a key player in putrescine-induced GABA production can prove to be fundamental to designing future studies that more deeply look into the role of this enzyme in astrocytes. The production of H_2_O_2_ and ammonia during the MAOB-mediated production of GABA are key molecules causing reactive astrogliosis in neurodegenerative diseases (Chun et al., 2020; Ju et al., 2022). While inhibition of ALDH1A1 or SIRT2, enzymes downstream to MAOB, would not reduce the production of these toxic molecules, the revelation of these enzymes in the pathway unearths a chance to understand astrocytic putrescine-to-GABA conversion and the cascade of events underlying neurodegeneration at the molecular level. The knowledge gained from this study will prove beneficial in gaining a deeper understanding of GABA production and provide new directions of study that were previously unexplored.

## ACKNOWLEDGEMENTS

This work was supported by the Institute for Basic Science (IBS), Center for Cognition and Sociality (IBS-R001-D2). This study was also supported by the National Research Foundation (NRF) Grants from the Korean Ministry of Education, Science and Technology (2018M3C7A1056894, NRF-2020M3E5D9079742) and KIST Grants (2E30954 and 2E30962).

## REFERENCES

1. Chai, H., Diaz-Castro, B., Shigetomi, E., Monte, E., Octeau, J.C., Yu, X., Cohn, W., Rajendran, P.S., Vondriska, T.M., and Whitelegge, J.P. (2017). Neural circuit-specialized astrocytes: transcriptomic, proteomic, morphological, and functional evidence. Neuron 95, 531–549. e539.

2. Chen, Y., Qin, C., Huang, J., Tang, X., Liu, C., Huang, K., Xu, J., Guo, G., Tong, A., and Zhou, L. (2020). The role of astrocytes in oxidative stress of central nervous system: A mixed blessing. Cell proliferation 53, e12781.

3. Chun, H., Im, H., Kang, Y.J., Kim, Y., Shin, J.H., Won, W., Lim, J., Ju, Y., Park, Y.M., Kim, S., et al. (2020). Severe reactive astrocytes precipitate pathological hallmarks of Alzheimer’s disease via H(2)O(2)(-) production. Nat Neurosci 23, 1555–1566.

4. Edenberg, H.J. (2007). The genetics of alcohol metabolism: role of alcohol dehydrogenase and aldehyde dehydrogenase variants. Alcohol Research & Health 30, 5.

5. Grabowska, W., Sikora, E., and Bielak-Zmijewska, A. (2017). Sirtuins, a promising target in slowing down the ageing process. Biogerontology 18, 447–476.

6. Heo, J.Y., Nam, M.-H., Yoon, H.H., Kim, J., Hwang, Y.J., Won, W., Woo, D.H., Lee, J.A., Park, H.-J., and Jo, S. (2020). Aberrant tonic inhibition of dopaminergic neuronal activity causes motor symptoms in animal models of Parkinson’s disease. Current Biology 30, 276–291. e279.

7. Ishibashi, M., Egawa, K., and Fukuda, A. (2019). Diverse actions of astrocytes in GABAergic signaling. International journal of molecular sciences 20, 2964.

8. Jin, S., Cao, Q., Yang, F., Zhu, H., Xu, S., Chen, Q., Wang, Z., Lin, Y., Cinar, R., and Pawlosky, R.J. (2021). Brain ethanol metabolism by astrocytic ALDH2 drives the behavioural effects of ethanol intoxication. Nature metabolism 3, 337–351.

9. Jo, S., Yarishkin, O., Hwang, Y.J., Chun, Y.E., Park, M., Woo, D.H., Bae, J.Y., Kim, T., Lee, J., and Chun, H. (2014). GABA from reactive astrocytes impairs memory in mouse models of Alzheimer’s disease. Nature medicine 20, 886–896.

10. Ju, Y.H., Bhalla, M., Hyeon, S.J., Oh, J.E., Yoo, S., Chae, U., Kwon, J., Koh, W., Lim, J., and Park, Y.M. (2022). Astrocytic urea cycle detoxifies Aβ-derived ammonia while impairing memory in Alzheimer’s disease. Cell Metabolism 34, 1104–1120. e1108.

11. Keung, W.M., and Vallee, B.L. (1998). Daidzin and its antidipsotropic analogs inhibit serotonin and dopamine metabolism in isolated mitochondria. Proc Natl Acad Sci U S A 95, 2198–2203.

12. Kim, J.-I., Ganesan, S., Luo, S.X., Wu, Y.-W., Park, E., Huang, E.J., Chen, L., and Ding, J.B. (2015). Aldehyde dehydrogenase 1a1 mediates a GABA synthesis pathway in midbrain dopaminergic neurons. Science 350, 102–106.

13. Kwak, H., Koh, W., Kim, S., Song, K., Shin, J.-I., Lee, J.M., Lee, E.H., Bae, J.Y., Ha, G.E., and Oh, J.-E. (2020). Astrocytes control sensory acuity via tonic inhibition in the thalamus. Neuron 108, 691–706. e610.

14. Le-Corronc, H., Rigo, J.-M., Branchereau, P., and Legendre, P. (2011). GABAA receptor and glycine receptor activation by paracrine/autocrine release of endogenous agonists: more than a simple communication pathway. Molecular neurobiology 44, 28–52.

15. Lee, M., Schwab, C., and Mcgeer, P.L. (2011). Astrocytes are GABAergic cells that modulate microglial activity. Glia 59, 152–165.

16. Li, W., Zhang, B., Tang, J., Cao, Q., Wu, Y., Wu, C., Guo, J., Ling, E.-A., and Liang, F. (2007). Sirtuin 2, a mammalian homolog of yeast silent information regulator-2 longevity regulator, is an oligodendroglial protein that decelerates cell differentiation through deacetylating α-tubulin. Journal of Neuroscience 27, 2606–2616.

17. Maxwell, M.M., Tomkinson, E.M., Nobles, J., Wizeman, J.W., Amore, A.M., Quinti, L., Chopra, V., Hersch, S.M., and Kazantsev, A.G. (2011). The Sirtuin 2 microtubule deacetylase is an abundant neuronal protein that accumulates in the aging CNS. Human molecular genetics 20, 3986–3996.

18. Morgan, C.A., Parajuli, B., Buchman, C.D., Dria, K., and Hurley, T.D. (2015). N, N-diethylaminobenzaldehyde (DEAB) as a substrate and mechanism-based inhibitor for human ALDH isoenzymes. Chemico-biological interactions 234, 18–28.

19. Nam, M.-H., Cho, J., Kwon, D.-H., Park, J.-Y., Woo, J., Lee, J.M., Lee, S., Ko, H.Y., Won, W., and Kim, R.G. (2020). Excessive astrocytic GABA causes cortical hypometabolism and impedes functional recovery after subcortical stroke. Cell reports 32, 107861.

20. Oh, S.-J., Lee, J.M., Kim, H.-B., Lee, J., Han, S., Bae, J.Y., Hong, G.-S., Koh, W., Kwon, J., and Hwang, E.-S. (2020). Ultrasonic neuromodulation via astrocytic TRPA1. Current Biology 30, 948.

21. Outeiro, T.F., Kontopoulos, E., Altmann, S.M., Kufareva, I., Strathearn, K.E., Amore, A.M., Volk, C.B., Maxwell, M.M., Rochet, J.-C., and McLean, P.J. (2007). Sirtuin 2 inhibitors rescue α-synuclein-mediated toxicity in models of Parkinson’s disease. science 317, 516–519.

22. Park, J.-H., Ju, Y.H., Choi, J.W., Song, H.J., Jang, B.K., Woo, J., Chun, H., Kim, H.J., Shin, S.J., and Yarishkin, O. (2019). Newly developed reversible MAO-B inhibitor circumvents the shortcomings of irreversible inhibitors in Alzheimer’s disease. Science advances 5, eaav0316.

23. Scuderi, C., Stecca, C., Bronzuoli, M.R., Rotili, D., Valente, S., Mai, A., and Steardo, L. (2014). Sirtuin modulators control reactive gliosis in an in vitro model of Alzheimer’s disease. Frontiers in Pharmacology 5, 89.

24. Singh, S., Kumar, P.U., Thakur, S., Kiran, S., Sen, B., Sharma, S., Rao, V.V., Poongothai, A., and Ramakrishna, G. (2015). Expression/localization patterns of sirtuins (SIRT1, SIRT2, and SIRT7) during progression of cervical cancer and effects of sirtuin inhibitors on growth of cervical cancer cells. Tumor Biology 36, 6159–6171.

25. Vaquero, A., Scher, M.B., Lee, D.H., Sutton, A., Cheng, H.-L., Alt, F.W., Serrano, L., Sternglanz, R., and Reinberg, D. (2006). SirT2 is a histone deacetylase with preference for histone H4 Lys 16 during mitosis. Genes & development 20, 1256–1261.

26. Wang, Y., Yang, J., Hong, T., Chen, X., and Cui, L. (2019). SIRT2: Controversy and multiple roles in disease and physiology. Ageing research reviews 55, 100961.

27. Woo, D.H., Han, K.-S., Shim, J.W., Yoon, B.-E., Kim, E., Bae, J.Y., Oh, S.-J., Hwang, E.M., Marmorstein, A.D., and Bae, Y.C. (2012). TREK-1 and Best1 channels mediate fast and slow glutamate release in astrocytes upon GPCR activation. Cell 151, 25–40.

28. Woo, J., Min, J.O., Kang, D.-S., Kim, Y.S., Jung, G.H., Park, H.J., Kim, S., An, H., Kwon, J., and Kim, J. (2018). Control of motor coordination by astrocytic tonic GABA release through modulation of excitation/inhibition balance in cerebellum. Proceedings of the National Academy of Sciences 115, 5004–5009.

29. Yeong, K.Y., Berdigaliyev, N., and Chang, Y. (2020). Sirtuins and their implications in neurodegenerative diseases from a drug discovery perspective. ACS chemical neuroscience 11, 4073–4091.

30. Yoon, B.-E., and Lee, C.J. (2014). GABA as a rising gliotransmitter. Frontiers in neural circuits 8, 141.

31. Yoon, B.E., Woo, J., Chun, Y.E., Chun, H., Jo, S., Bae, J.Y., An, H., Min, J.O., Oh, S.J., and Han, K.S. (2014). Glial GABA, synthesized by monoamine oxidase B, mediates tonic inhibition. The Journal of physiology 592, 4951–4968.

